# Domain Acquisition by Class I Aminoacyl-tRNA Synthetase Urzymes Coordinated the Catalytic Functions of HVGH and KMSKS Motifs

**DOI:** 10.1101/2023.04.26.538376

**Authors:** Guo Qing Tang, Jessica J. Hobson Elder, Jordan Douglas, Charles W. Carter

## Abstract

Leucyl-tRNA synthetase (LeuRS) is a Class I aminoacyl-tRNA synthetase (aaRS) that catalyzes synthesis of leucyl-tRNA^leu^ for codon-directed protein synthesis on the ribosome. Class I aaRS, which were key to the evolution of genetic coding, contain two discrete signature sequences, HIGH and KMSKS, that participate in transition-state stabilization by the entire eleven-enzyme Class I aaRS superfamily. Combinatorial mutagenesis and thermodynamic cycle analyses of these catalytic signatures in full-length *Pyrococcus horikoshii* LeuRS and the 129-residue urzyme ancestral model generated from it (LeuAC) provide quantitative insight into the evolutionary gain of function induced by acquisition of the anticodon-binding (ABD) and multiple insertion modules in the catalytic domain. The free energy coupling terms, Δ(ΔG^‡^), are small and unfavorable for LeuAC, but large and favorable for LeuRS. Thus, the ABD and other insertion modules induce strong cooperativity between the two signature sequences, which are uncoupled in LeuAC. These results further substantiate the authenticity of LeuAC urzyme catalysis and implicate domain motion in catalysis by the full-length LeuRS. Most importantly, the implication that backbone elements of secondary structures achieve a major portion of the overall transition-state stabilization by LeuAC is also consistent with coevolution of the genetic code and metabolic pathways necessary to produce histidine and lysine sidechains.

**Bullet Points:**

1. The LeuRS HVGH and KMSKS signature motifs are energetically coupled by −1.6 kcal/mole.
2. The same motifs are anti-coupled by +0.8 kcal/mole in the 129 residue urzyme, LeuAC.
3. Ancestral Class I aaRS did not require either histidine or lysine for catalysis.

## 1. INTRODUCTION

Class I aminoacyl-tRNA synthetases (aaRS) are one of two superfamilies that, together with their cognate tRNAs, translate the genetic code(1). They perform that function by activating cognate amino acids at the expense of the two high-energy bonds in ATP and transferring the activated aminoacyl group from the 5’ position of AMP to the tRNA 3’-terminal ribose. Eleven contemporary Class I aaRS families are responsible for the hydrophobic amino acids (isoleucine, valine, leucine, methionine), together with the larger homologs of the acidic (glutamate), amide (glutamine), and aromatic (tryptophan, tyrosine) amino acids, as well as arginine and cysteine. Some archaea and bacteria also possess a Class I lysyl-tRNA synthetase(2), which in most organisms is a Class II enzyme. All Class I aaRS share the same three domains: a Rossmann dinucleotide-binding catalytic domain with variable length insertions between the two crossover connections of the Rossmann fold (CP)(3,4) with an embedded editing domain in larger Class I enzymes, and an idiosyncratic anticodon-binding domain, ABD. The active site within the Rossmann fold is highly conserved across the superfamily.(5) Moreover, that domain has been re-constructed without any of the other domains, and it retains all three of the functions—amino acid activation, retention of the activated amino acid, and transfer to cognate tRNA—necessary to implement a rudimentary form of the code. For these reasons, we have argued that such constructs, called “urzymes”(6), are excellent experimental models for studying the emergence and early evolutionary history of ancestral aaRS(7-10) and genetic coding generally.

Here, we utilize the leucyl-tRNA synthetase from *Pyrococcus horikoshii*, LeuRS(11) and its urzyme, LeuAC(12), to examine the evolutionary gain of function associated with acquiring the CP and ABD domains. We note that *P. horikoshii* expresses the archaeal-like LeuRS and the location of its post-transfer editing domain within the extended connecting peptide (CP) insertion differs from that in eubacterial LeuRSs.(13) Construction of the LeuAC urzyme entails removing the entire extended CP domain, which includes many acquired insertion modules(14) as well as the ABD and C-terminal domains (Figure 1). LeuAC is thus homologous in that respect to the corresponding urzyme excerpted from *Geobacillus stearothermophilus*, TrpAC(6,15). However, the mass deleted from the catalytic domain (377 amino acids) is ∼ five times more than were deleted (74 amino acids) from the TrpRS catalytic domain. That missing mass includes insertion modules that others have suggested were essential to aminoacylation.(16)

**Figure 1.**
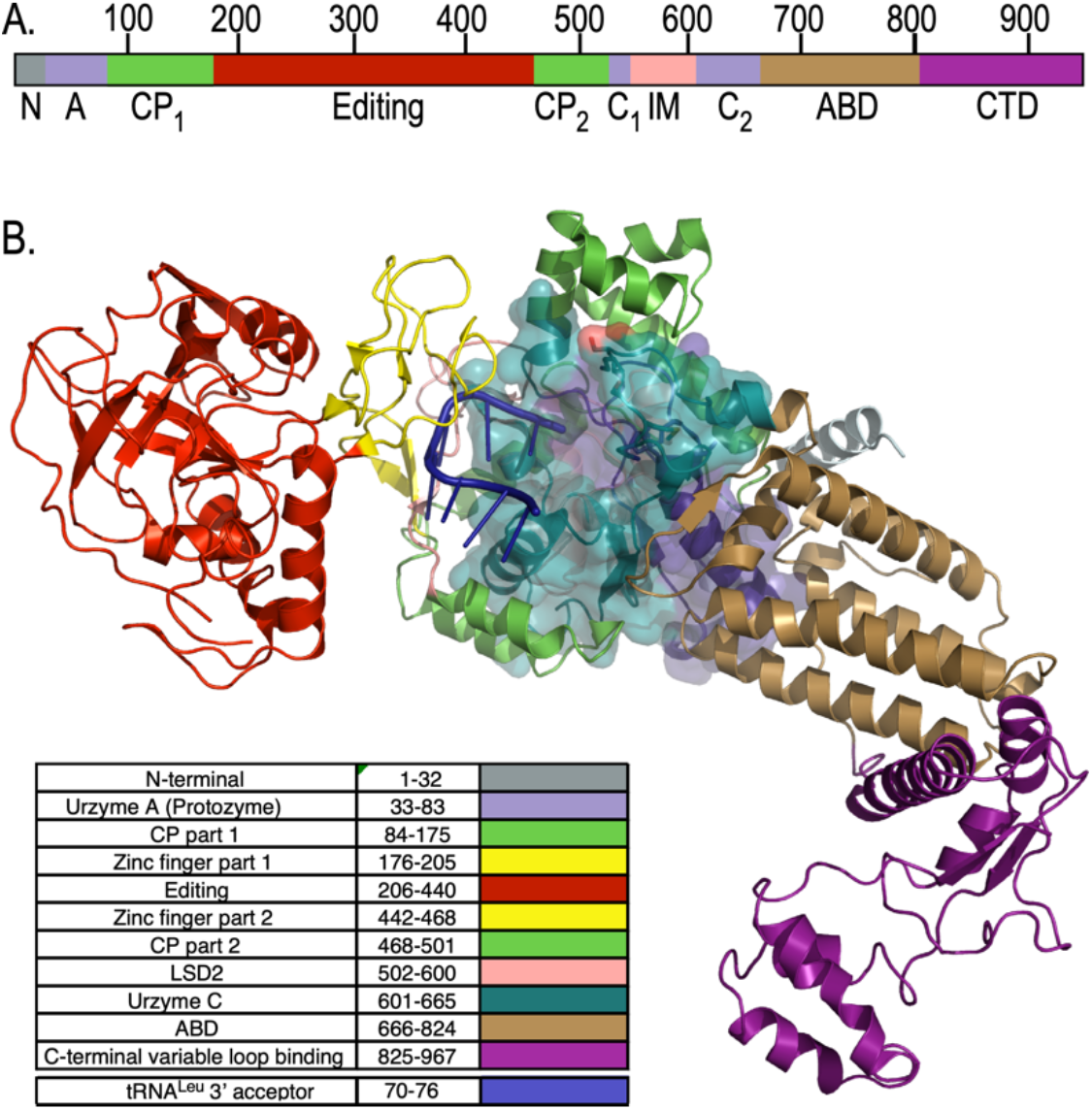
Relation of LeuAC urzyme to full-length *P. horikoshii* LeuRS. Colors are consistent between A and B. **A**. Linear schematic of LeuRS domains. N, N-terminal helical domain; A, Protozyme (ATP-binding site); CP, connecting peptide, a nested insertion containing the editing domain; Editing, Editing domain; C_1_, second fragment of the urzyme (amino acid and tRNA binding). LSD_2_, Leucine specific domain 2(11), an insertion module before second crossover; C_2_, C-terminal fragment of the urzyme (pyrophosphate binding); ABD, anticodon-binding domain; CTD, C-terminal domain that binds to the tRNA^Leu^ variable loop. **B**. Three dimensional cartoon based on PDB ID 1WC2 showing the LeuAC urzyme (surface embedded into the full-length enzyme and the arrangement of modules acquired during the specialization of LeuRS from other Class I aaRS. A76 of the tRNA^Leu^ acceptor stem inserts into the active sites of both LeuRS and LeuAC, bisecting the protozyme (blue) and C_2_, the second half of the urzyme. Insertions into a loop connecting the protozyme and C2 are all nested, one within the next, forming the complete CP. In contrast, C-terminal additions are serial.

**Figure 1.**
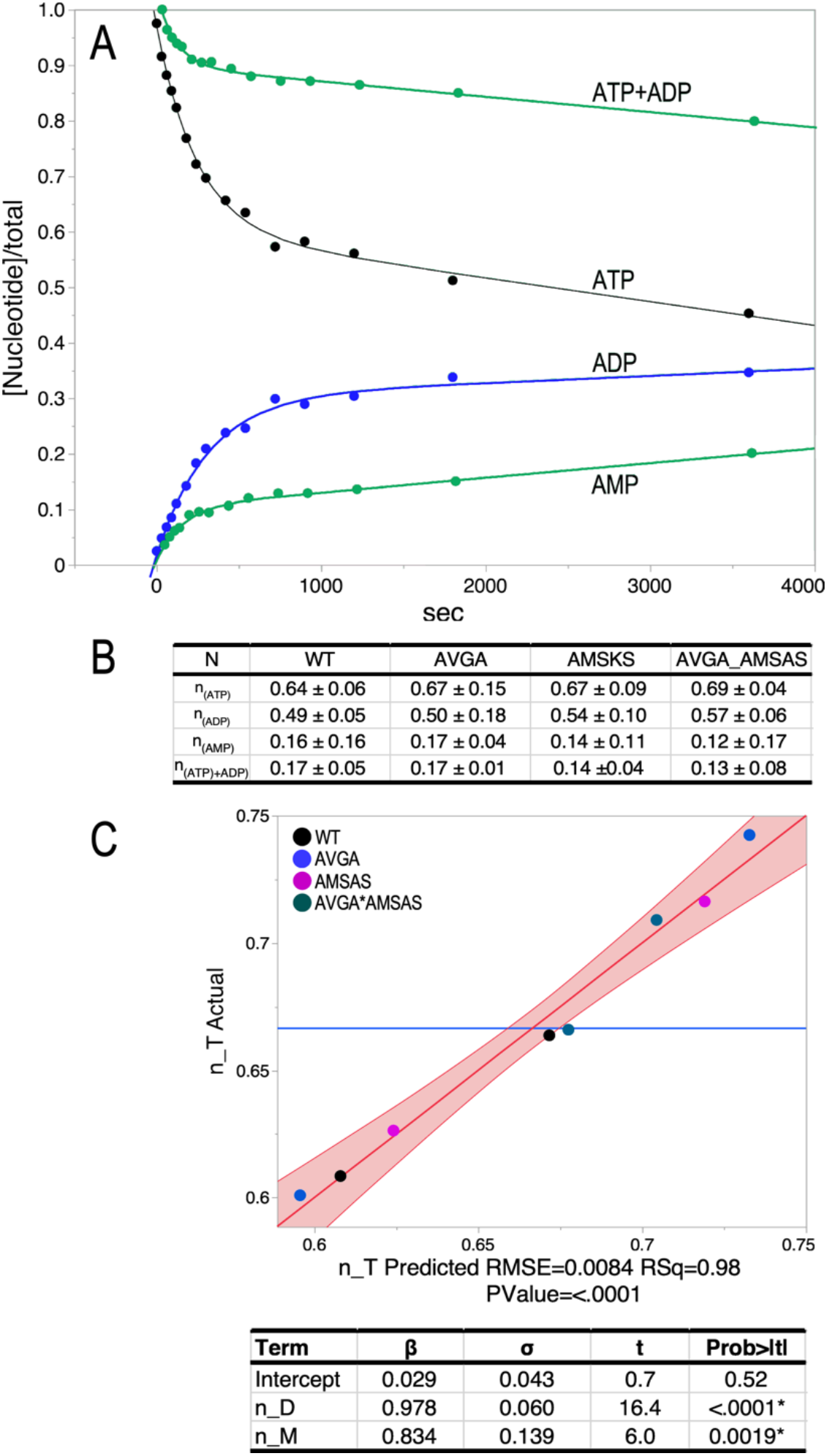
N-value determination for the four LeuAC variants. **A**. Single turnover time courses for the three adenine nucleotides. The green ATP + ADP curve is obtained by adding back the ATP converted to ADP (blue curve). It represents the ATP consumption after accounting for ADP production and is almost a reflection of the (green) curve for AMP production. Thus, it likely represents in situ formation of leu-5’AMP. **B**. Averaged n-values determined from the two physical constants, A (the amplitude of the burst) and C (the offset), with their standard deviations obtained for each variant by fitting to Eqn. (1) as described in Methods and Materials. **C**. Multivariate regression analysis of n-values averaged in **B** confirm that n-values for the appearance of ADP and AMP account almost exactly for ATP consumption. Solving the simultaneous equations [n_T_i_ = Σ_i,variants_ (β_D_(C + n_D) + β_M_(n_M) + ε)] by least squares using JMP16PRO(31) gave the table of values underneath the graph. Here, and elsewhere, the true value of the regression line has a 95 % chance of falling within the red boundaries.

Two consensus sequence motifs present in the Rossmann fold (HIGH and KMSKS) have been implicated multiple ways in catalysis of both amino acid activation and acyl-transfer to tRNA in many Class I aaRS. Their conservation across the superfamily has been an evolutionary curiosity, suggesting that they are central to catalysis. However, both histidine and lysine require complex biosynthetic pathways, and thus are unlikely to have been available for the earliest stages of codon-dependent translation.(17-20)

The unexpectedly modest impact of mutating lysine residues to alanine in the LeuAC urzyme KMSKS signature(12) motivated us to examine the catalytic roles of both signatures in both LeuRS and LeuAC. An obvious way to do this is with a factorial design in which both histidines in the HVGH signature are also mutated to alanine separately and in combination. To our knowledge, and despite a landmark combinatorial mutagenesis of the KFGKT sequence in TyrRS(21-24), no one has examined the coordinated behavior of the two Class I catalytic signatures, even in a full-length Class I aaRS. That combinatorial mutagenesis facilitates measurement of the energetic coupling between the two signatures. Comparing that coupling in urzyme and full-length enzyme also maps changes in the evolutionary time domain.

The comparison reinforces previous suggestions(25-28) that full-length Class I aaRS benefit from a significant intramolecular cooperativity that is absent from the corresponding urzymes. We identify side chain packing interactions between the hydrophobic side chains in each catalytic signature and a cluster of nonpolar side chains in the anticodon-binding domain that is conserved throughout the Class. Embedding the valine and methionine side chains of the LeuRS HVGH and KMSKS sequences into that cluster provides a structural rationale for their coordination during the catalytic cycle, consistent with their catalytic coupling in the full-length enzyme.

The anti-coupling between HVGH and KMSKS signatures in the urzyme has important implications for the emergence and early evolution of polypeptide catalysts. Surprisingly, the AVGA variant of the LeuAC urzyme benefits from both the wild type KMSKS signature and unfavorable coupling energy; it is thus the most active LeuAC variant and is actually more active at aminoacylation than the corresponding mutant of full-length LeuRS. Catalysis by aaRS urzymes does not require specific amino acid side chains that are highly conserved in the active sites of all Class I aaRS, thus significantly broadening the sequence space for primordial catalysts and creating substantive experimental support for Wong’s proposal that the genetic code coevolved with the amino acid biosynthetic pathways.(17,18)

Finally, the systematic consistency of thermodynamic cycle analysis and provision of a structural rationale for the differential intramolecular coupling in the full-length enzyme also decisively confirm the authenticity of catalysis by aaRS urzymes.

## 2. RESULTS

The experimental design matrix for this work is the 2^3^ factorial design in which all possible combinations are made of HVGH → AVGA and KMSKS → AMSAS mutations in both full-length LeuRS and the LeuAC urzyme. The eight variants were constructed, expressed, and purified as described previously(12) and in Materials and Methods. We assayed all variants for leucine activation by single turnover active-site titrations (Table 1) and for aminoacylation of tRNA^Leu^ in triplicate (Table 3).

**Table 1.** Design matrix for single-turnover kinetic assays to determine the fraction (n) of active molecules. The first four columns are the estimated parameters from Eqn. (1). Columns 11-17 are binary entries specifying independent variables.

|  | C | A | k<sub>chem</sub> | k<sub>cat</sub> | k<sub>chem</sub>/k<sub>cat</sub> | ΔG<sub>kchem</sub> | ΔG<sub>kcat</sub> | ΔG<sub>kchem</sub>/k<sub>cat</sub> | ATP | ADP | AMP |
| --- | --- | --- | --- | --- | --- | --- | --- | --- | --- | --- | --- |
| **LeuAC** |  |  |  |  |  |  |  |  |  |  |  |
| **ATP** |  |  |  |  |  |  |  |  |  |  |  |
| WT | 0.60 | 0.36 | 0.004 | 0.000043 | 96 | 3.26 | 5.96 | -2.70 | 1 | 0 | 0 |
| AVGA | 0.55 | 0.36 | 0.015 | 0.000043 | 342 | 2.49 | 5.95 | -3.45 | 1 | 0 | 0 |
| AMSAS | 0.57 | 0.38 | 0.005 | 0.000044 | 106 | 3.18 | 5.94 | -2.76 | 1 | 0 | 0 |
| AVGA_AMSAS | 0.57 | 0.40 | 0.008 | 0.000023 | 321 | 2.89 | 6.31 | -3.42 | 1 | 0 | 0 |
| **ADP** |  |  |  |  |  |  |  |  |  |  |  |
| WT | 0.30 | 0.28 | 0.003 | 0.000013 | 252 | 3.40 | 6.67 | -3.27 | 0 | 1 | 0 |
| AVGA | 0.34 | 0.26 | 0.019 | 0.000036 | 518 | 2.36 | 6.06 | -3.70 | 0 | 1 | 0 |
| AMSAS | 0.35 | 0.30 | 0.004 | 0.000013 | 319 | 3.27 | 6.68 | -3.41 | 0 | 1 | 0 |
| AVGA_AMSAS | 0.36 | 0.33 | 0.008 | 0.000011 | 706 | 2.86 | 6.74 | -3.88 | 0 | 1 | 0 |
| **AMP** |  |  |  |  |  |  |  |  |  |  |  |
| WT | 0.11 | 0.08 | 0.006 | 0.000026 | 243 | 2.99 | 6.24 | -3.25 | 0 | 0 | 1 |
| AVGA | 0.10 | 0.10 | 0.009 | 0.000007 | 1266 | 2.80 | 7.03 | -4.23 | 0 | 0 | 1 |
| AMSAS | 0.09 | 0.08 | 0.006 | 0.000029 | 221 | 2.98 | 6.18 | -3.20 | 0 | 0 | 1 |
| AVGA_AMSAS | 0.06 | 0.08 | 0.007 | 0.000013 | 577 | 2.90 | 6.67 | -3.76 | 0 | 0 | 1 |
| **ATP+ADP** |  |  |  |  |  |  |  |  |  |  |  |
| WT | 0.90 | 0.10 | 0.01 | 2.8E-05 | 311 | 2.82 | 6.22 | -3.40 | 0 | 0 | 0 |
| AVGA | 0.90 | 0.10 | 0.01 | 7E-06 | 1305 | 2.77 | 7.01 | -4.25 | 0 | 0 | 0 |
| AMSAS | 0.91 | 0.08 | 0.01 | 0.00003 | 256 | 2.89 | 6.17 | -3.28 | 0 | 0 | 0 |
| AVGA_AMSAS | 0.93 | 0.08 | 0 | 3E-06 | 1274 | 3.23 | 7.47 | -4.23 | 0 | 0 | 0 |

### 2.1 Single turnover experiments give consistent first-order rates for changes in ATP, ADP, and AMP

Single turnover kinetic experiments, often called active-site titrations (AST) are done using substrate level amounts of enzyme. They measure the size of the burst corresponding to the first round of catalysis and are therefore the method of choice to estimate fractional activities of enzyme variants(29,30) and normalizing enzyme concentrations when computing k_cat_ values. AST assays performed to estimate the active fractions in all variants provide supplemental evidence that mutational impacts are comparable in both leucine activation and aminoacylation by LeuAC.

We uncovered an unexpected phenomenon when we followed the time courses for all three adenine nucleotides in AST assays of aaRS urzymes (Fig. 1). ATP consumption is accompanied by ADP production(12), in addition to the expected products, aminoacyl-5’AMP and AMP, which is produced by product release and hydrolysis. Prior to analyzing kinetic data for the thermodynamic cycle, we studied these curves in detail to assure a consistent interpretation.

Quantitation of the active-site titration curves (Fig. 1A) confirms first that they are additive; n-values (Fig. 1B) from ATP consumption are quantitative sums of those for ADP and AMP produced (Fig. 1C). LeuAC variant n-values from fitting C and A in Eqn. (1) agree with a fractional error of ∼ 0.09. ATP consumption <n_T> values range from 0.64 – 0.69 for the different combinatorial variants. <n_D> values (ADP) range from 0.49 – 0.57. <n_M> values (AMP) range from 0.13 – 0.17. In each case n_T = n_D + n_M within 2 % (i.e., intercepts are ∼0; slopes are both ∼1). These data furnish a self-consistent picture of multiple catalyzed reactions occurring in the first round of catalysis (Table 2).

**Table 2.** Relative activities of LeuAC mutants in the first round of catalysis. Values in the table are taken from n-values determined by active-site titration.

|  | WT | AVGA | AMSAS | AVGA*AMSAS |
| --- | --- | --- | --- | --- |
| Active fraction | 0.64 ±0.04 | 0.67 ± 0.10 | 0.67 ± 0.06 | 0.69 ± 0.03 |
| ADP/Tot | 0.49 | 0.50 | 0.54 | 0.57 |
| AMP/Tot | 0.16 | 0.17 | 0.14 | 0.13 |
| ADP % | 0.77 | 0.75 | 0.80 | 0.83 |
| AMP % | 0.25 | 0.25 | 0.21 | 0.18 |

**Table 3.** Design matrix with aminoacylation rates and transition-state stabilization free energies for the eight LeuRS variants, together with codes for the three independent variables.

| Catalyst | k(/mole/sec<sup>a</sup>) | $\Delta G^\ddagger$(kcal/mole) | KMSKS | HVGH | HVGA*KMSKS | FULL |
| --- | --- | --- | --- | --- | --- | --- |
| LeuAC_WT | 0.0085 | 2.82 | 1 | 1 | 1 | 0 |
| LeuAC_WT | 0.0081 | 2.85 | 1 | 1 | 1 | 0 |
| LeuAC_WT | 0.0056 | 2.78 | 1 | 1 | 1 | 0 |
| LeuAC_WT | 0.0067 | 2.67 | 1 | 1 | 1 | 0 |
| LeuAC_WT | 0.0051 | 2.83 | 1 | 1 | 1 | 0 |
| LeuAC_AVGA | 0.0193 | 2.04 | 1 | 0 | 0 | 0 |
| LeuAC_AVGA | 0.0339 | 1.71 | 1 | 0 | 0 | 0 |
| LeuAC_AVGA | 0.0358 | 1.68 | 1 | 0 | 0 | 0 |
| LeuAC_AVGA | 0.0390 | 1.92 | 1 | 0 | 0 | 0 |
| LeuAC_AVGA | 0.0383 | 1.93 | 1 | 0 | 0 | 0 |
| LeuAC_AMSAS | 0.0063 | 3.00 | 0 | 1 | 0 | 0 |
| LeuAC_AMSAS | 0.0065 | 2.98 | 0 | 1 | 0 | 0 |
| LeuAC_AMSAS | 0.0088 | 2.80 | 0 | 1 | 0 | 0 |
| LeuAC_AMSAS | 0.0067 | 2.67 | 0 | 1 | 0 | 0 |
| LeuAC_AMSAS | 0.0057 | 2.77 | 0 | 1 | 0 | 0 |
| LeuAC_AMSAS | 0.0068 | 2.66 | 0 | 1 | 0 | 0 |
| LeuAC_AVGA+AMSAS | 0.0188 | 2.35 | 0 | 0 | 0 | 0 |
| LeuAC_AVGA+AMSAS | 0.0057 | 3.06 | 0 | 0 | 0 | 0 |
| LeuAC_AVGA+AMSAS | 0.0067 | 2.97 | 0 | 0 | 0 | 0 |
| LeuAC_AVGA+AMSAS | 0.0119 | 2.33 | 0 | 0 | 0 | 0 |
| LeuAC_AVGA+AMSAS | 0.0053 | 2.81 | 0 | 0 | 0 | 0 |
| LeuAC_AVGA+AMSAS | 0.0079 | 2.57 | 0 | 0 | 0 | 0 |
| FL-LeuRS_WT | 0.3216 | 0.71 | 1 | 1 | 1 | 1 |
| FL-LeuRS_WT | 0.3173 | 0.72 | 1 | 1 | 1 | 1 |
| FL-LeuRS_WT | 0.3011 | 0.75 | 1 | 1 | 1 | 1 |
| FL-LeuRS_AVGA | 0.0168 | 2.58 | 1 | 0 | 0 | 1 |
| FL-LeuRS_AVGA | 0.0423 | 2.03 | 1 | 0 | 0 | 1 |
| FL-LeuRS_AVGA | 0.0811 | 1.65 | 1 | 0 | 0 | 1 |
| FL-LeuRS_AMSAS | 0.0872 | 1.64 | 0 | 1 | 0 | 1 |
| FL-LeuRS_AMSAS | 0.1385 | 1.37 | 0 | 1 | 0 | 1 |
| FL-LeuRS_AMSAS | 0.1549 | 1.30 | 0 | 1 | 0 | 1 |
| FL-LeuRS_AVGA+AMSAS | 0.4131 | 1.03 | 0 | 0 | 0 | 1 |
| FL-LeuRS_AVGA+AMSAS | 0.2465 | 1.33 | 0 | 0 | 0 | 1 |
| FL-LeuRS_AVGA+AMSAS | 0.3175 | 1.18 | 0 | 0 | 0 | 1 |

<sup>a</sup> Acylation rates have been corrected by dividing by both fractional activity of the respective catalyst and the fractional activity of substrate tRNA.

LeuAC variant-specific column vectors in Fig. 1B are quite strictly parallel; all six correlation coefficients have R^2^ values >0.99. Thus, although combinatorial mutagenesis changes the active fractions, it has no effect on the relative magnitudes of the four different n-values. Strict linear dependence means that, for the purpose of normalizing the active fractions of different variants in analyzing aminoacylation rates, any of the n-values would be equally suitable. For the present purpose, we chose to normalize acylation rates using the n_T values, to estimate the minimum rate enhancements of acylation by full length LeuRS, relative to LeuAC. Although the burst sizes are well-determined in AST assays of LeuRS, the rapid time scale precludes sufficiently precise determination of rate constants to estimate the effects of combinatorial mutagenesis. Thus, we did not attempt to analyze them.

### 2.2 Combinatorial mutants of HIGH and KMSKS catalytic signatures in LeuRS reduce its catalytic proficiency for tRNA^Leu^ acylation to that of LeuAC and its mutational variants

A histogram of transition-state stabilization free energies for tRNA^Leu^ aminoacylation, ΔG^‡^, for all eight variants (Fig. 2) reveals that the two catalytic signatures make profound contributions to the catalytic proficiency of the full-length enzyme. The distributions in the upper right of Fig. 2B illustrate that ΔG^‡^ measurements for the mutations in full-length LeuRS have values comparable to those of the wild type Urzyme LeuAC and its corresponding mutations. Rather than forming a separate distribution in the neighborhood of the WT LeuRS, as observed for LeuAC variants, all LeuRS mutants are about as compromised as the LeuAC urzyme and its combinatorial mutants. Active-site mutations therefore have severe effects on the full-length enzyme, but substantially smaller effects on the urzyme. The wild-type urzyme active site therefore cannot achieve a comparable rate enhancement chemistry to that of the full-length enzyme.

**Figure 2.**
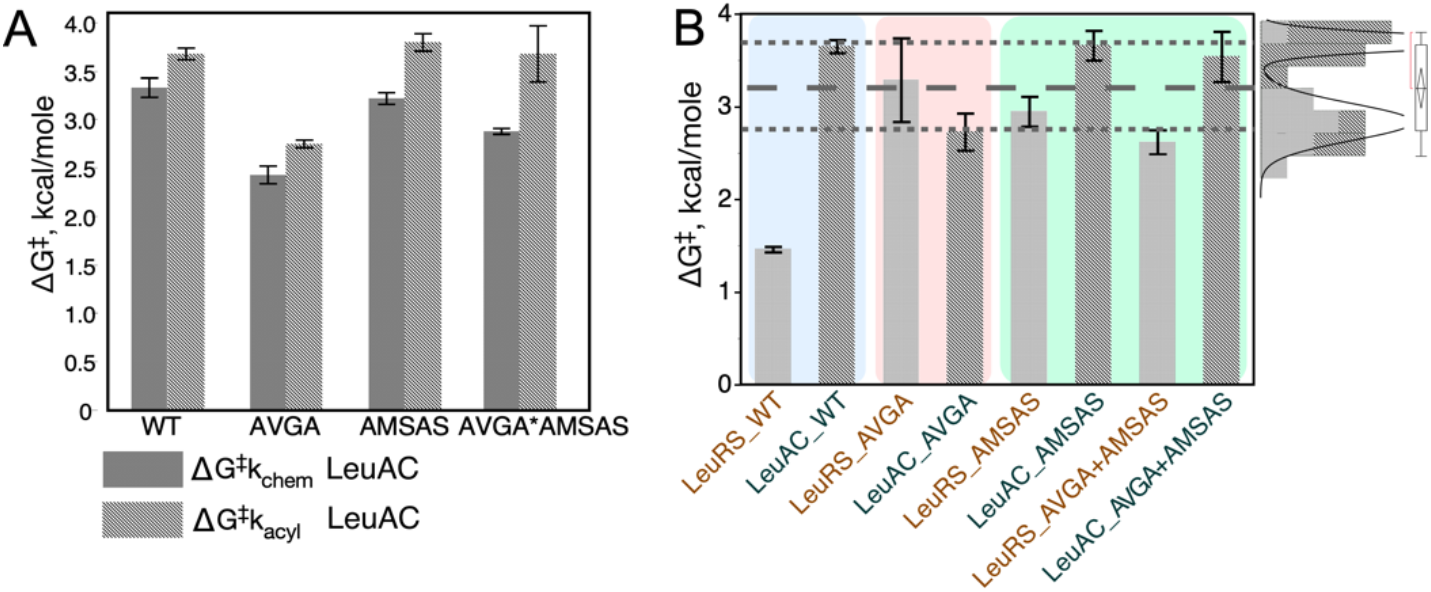
Individual transition-state stabilization free energies, ΔG^‡^, in kcal/mole for each LeuRS variant identified along the x axis. Error bars are two-way 95 % confidence levels from triplicate determinations. **A**. Rate accelerations for all LeuAC variants show comparable changes in activity in amino acid activation and tRNA^Leu^ aminoacylation (§2.3). They are greater for activation than for acylation, as expected from differences in the corresponding uncatalyzed rates(32). **B**. LeuRS variants are generally stronger catalysts than LeuAC variants, with the largest gap being between variants with wild type sequences (blue panel). AVGA variants have inverted rates, (rose panel) because the AVGA LeuAC variant has greater activity than the corresponding variant of full-length LeuRS. Finally, AMSAS and double mutant variants have reduced activity in LeuAC and increased activity in LeuRS, relative to the corresponding AVGA variants (green panel). Panel in the upper right shows the distribution of all 21 measurements for variants excepting WT LeuRS. Grey dashed lines identify a significant subset whose mean ΔG^‡^, ∼3.2 kcal/mole, is significantly greater than that of WT LeuRS. Mutational variants of LeuRS, excepting AVGA, are nonetheless also better catalysts than those of corresponding LeuAC. This significant variation is difficult to interpret without thermodynamic cycle analysis (§2.4).

### 2.3 Combinatorial mutagenesis has comparable impacts on leucine activation and tRNA^Leu^ aminoacylation by LeuAC

AST assays (Fig. 1) furnish values of the internal first-order rate constant for the first round catalysis of leucine activation. Using these values, we can compare the impacts of mutation on the activation reaction. Fig. 2A compares transition-state stabilization free energies of the four LeuAC variants in the two successive steps of tRNA^Leu^ biosynthesis. Qualitatively, the main difference between them is that all variants have lower ΔG^‡^ for activation than for aminoacylation, in keeping with their relative uncatalyzed rates. The AVGA mutant is markedly faster than any of the others, and the AMSAS variant is slowest by a small margin over the double mutant. These differences are easier to appreciate when transformed by estimating parameters for the corresponding thermodynamic cycle (§2.4).

### 2.4 Thermodynamic cycle analysis emphasizes the gain of function resulting from ABD and CP domain acquisition

The factorial design matrices (Tables I,II) enable us to estimate the magnitude and statistical significance of free energy contributions to transition-state stabilization that can be attributed to the intrinsic effects of each mutant group, as well as the magnitude of the synergy between them. For this purpose we use a construct known as a thermodynamic cycle, advocated by Jencks(33) and popularized by Horovitz and Fersht(34). As originally formulated, construction of a thermodynamic cycle entails computing differences observed in activation free energies, Δ(ΔG^‡^), at each corner of the polygon implicit in the factorial design. For a double mutant cycle, that involves a square with WT enzyme, the two single mutants, and the double mutant at the corners of a square.

We showed(27) that in practice it is more convenient to use multiple regression methods to estimate the coefficients in Eqn. (2). When values of the experimental error, ε, are small and the percentage of the variation in experimental data points that is explained by the regression model is high (≥ 0.9), a thermodynamic cycle is equivalent to changing the coordinate system in which the experimental free energies in Fig. 2 are presented.

The regression model for the thermodynamic cycle for ΔG^‡^_kchem_ from first-order rates, for all three nucleotides either consumed or produced in amino acid activation (Fig. 3) reveals several notable details. First, Studentized residuals (the prediction error, ε, divided by its adjusted standard error) associated with the contributing data points are distributed between –4 ≤ 4, so there are no outliers. Second, there are five significant predictors listed in the table. Their Student t-test probabilities are all ≤ 0.02, hence are statistically significant. Third, because neither ATP nor ADP are significant predictors of the first-order rate, there is no significant difference between rates for ATP consumption and ADP formation for the four variants. AMP production, on the other hand, is significantly different, and the AMP*HIGH β-coefficient means that it depends on whether the HIGH sequence is native or mutant. We return to this point after considering mutational effects on tRNA^Leu^ acylation.

**Figure 3.**
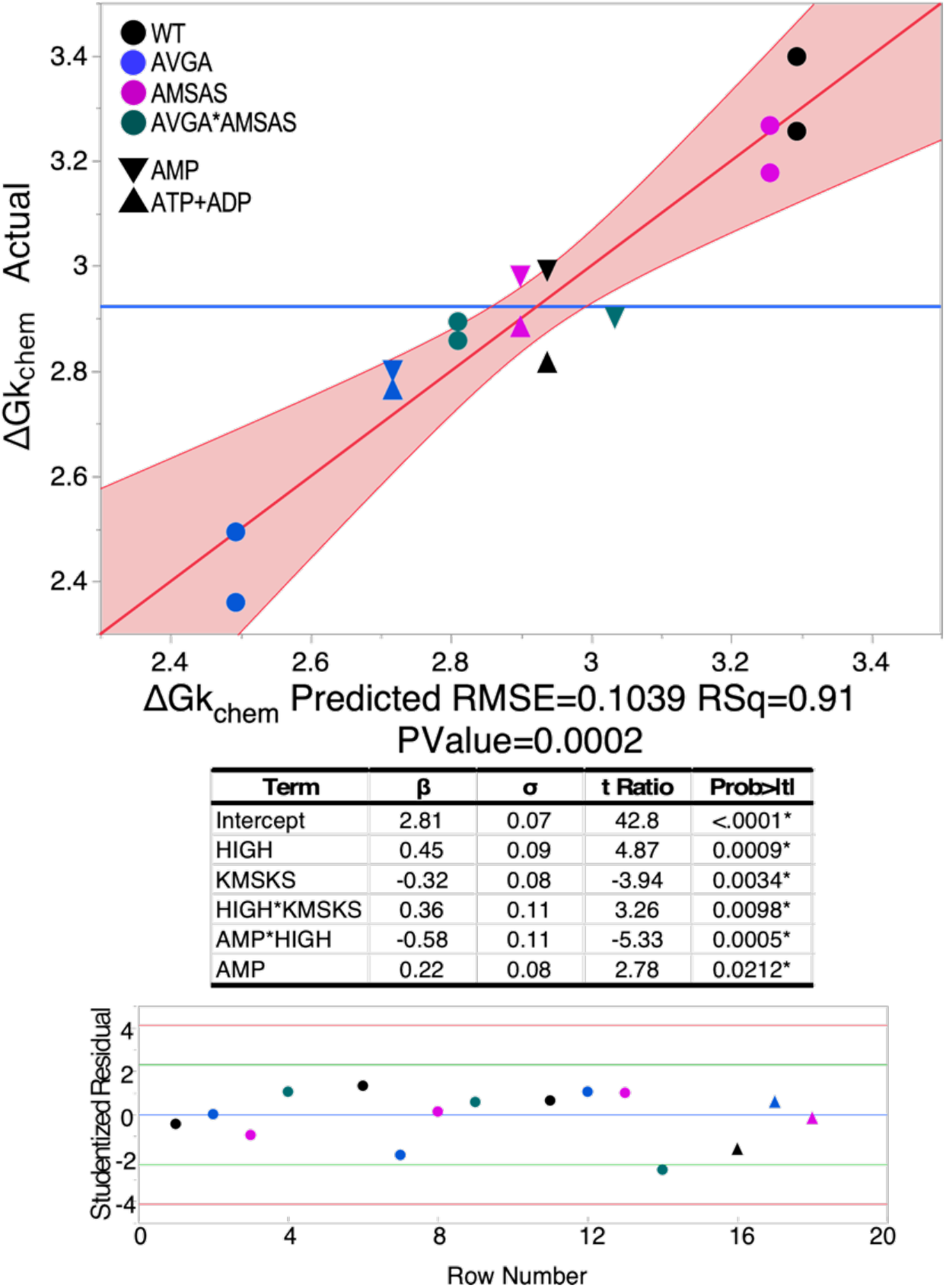
Multiple regression model for the activation free energies, ΔG^‡^k_chem_ for the internal first-order rate constant, k_chem_, for all LeuAC variants for time courses in Fig. 1A. Circles represent ATP consumption and ADP production; triangles represent the two alternative measurements for AMP production. β-coefficients are in kcal/mole. Variants are colored according to the scheme in the upper left-hand corner. Studentized residuals (ε_St_, bottom) are all –4.0< ε_St_ <4.0 confirming the absence of outliers.

First-order rate constants from AST, k_chem_, are independent of enzyme concentrations, and so do not require normalization for active fraction before converting to free energies for multiple regression analysis to construct thermodynamic cycles (Fig. 3,4). Fig. 3 highlights the statistical quality of the regression model; the corresponding models are not shown in Fig. 4. Fig. 4 displays histograms of the β-coefficients in kcal/mole with error bars denoting the 95% confidence levels. LeuAC and LeuRS histograms are placed within the respective thermodynamic cycles. Arrows form a cycle showing that the net free energy change of the successive steps total zero. The difference between left and right vertical arrows must equal that between top and bottom arrows, and is equal to the free energy, Δ(ΔG^‡^) for the HVGH*KMSKS interaction.

**Figure 4.**
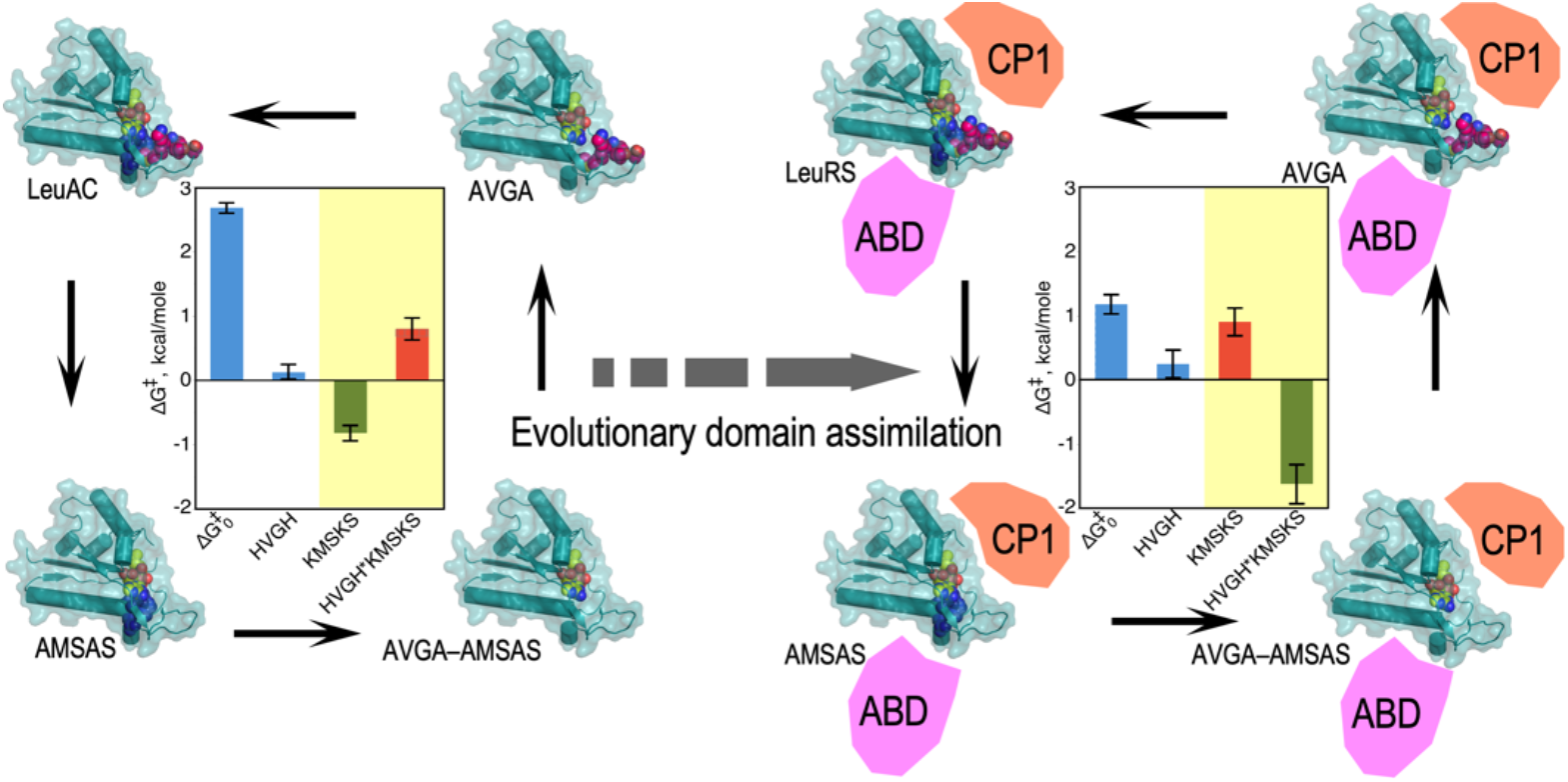
Thermodynamic cycles for LeuAC (left) and LeuRS. Transformation of data in Fig. 2B into its components via multiple regression analysis to determine successive changes in activation free energy, (ΔG^‡^), for successive mutations shown by single-headed arrows reveals that intrinsic effects of LeuAC and LeuRS catalysts and their HVGH signatures are similar in sign and relative magnitude (white panel). KMSKS and two-way HVGH*KMSKS Δ(ΔG^‡^) values however behave radically differently with and without the anticodon-binding (ABD) and CP domains (yellow panels). Specifically, the two-way synergy between the two signatures changes from anti-cooperative and unfavorable in LeuAC (red bar) to strongly cooperative and favorable (green bar) in LeuRS, leading to an increase of ∼2.6 kcal/mole in transition-state stabilization for aminoacylation of tRNA^Leu^. CP and ABD domains in full-length LeuRS are shown schematically.

The most striking feature in Fig. 3 is that samples at the extremes (the black circles = WT; the blue circles = AVGA) are opposite of what might be expected. The AVGA mutant is, by ∼ 1 kcal/mole, the most active catalyst in leucine activation and the WT variant is least active. Moreover, although the sign of the β_KMSKS_ coefficient is negative, signifying that the wild-type lysine residues favor catalysis, the synergy between it and the HIGH sequence, β_HVGH_*_KMSKS_, is positive. Thus, when it comes to stabilizing the transition state for the internally catalyzed reaction, the two signatures actually work against one another. That, and the positive β_HIGH_ value, account for the superiority of the AVGA variant, which is ∼5 times more active that wild-type LeuAC.

Finally, β coefficients for the main effects of the two mutations and their interaction tell us much about how the two catalytic signatures impact LeuAC catalysis. Wild-type residues in each location have opposite effects. Wild-type lysine residues in the KMSKS signature enhance catalysis modestly (ΔG^‡^_KMSKS_ = –0.32 kcal/mole) compared to alanine mutants. Wild-type histidine residues in the HVGH signature actually reduce catalysis by a comparable amount (positive ΔG^‡^_HVGH_ = 0.45 kcal/mole). The interaction term is also positive (ΔG^‡^_HVGH*KMSKS_ = 0.36 kcal/mole), so the negative impact of the native HVGH sequence is intensified by the wild-type lysine residues in the KMSKS signature, and seemingly paradoxically, increased catalysis by mutant AVGA sequence is intensified in the presence of wild type KMSKS.

Thermodynamic cycles for tRNA^Leu^ aminoacylation are similar, as suggested by Fig. 2A. Corresponding cycles for combinatorial mutagenesis in LeuAC and LeuRS are compared in Fig. 4. ΔG^‡^_0_ is the mean activation free energy for all four variants of the respective catalysts; the additional transition-state stabilization (β<0) or destabilization (β>0) attributable to wild type HVGH, KMSKS, sequences and their two-way interaction—or synergy—are represented by the remaining bars of the histograms. The inversion of the KMSKS and HVGH*KMSKS effects represents catalytic consequences of adding the anticodon binding domain and CP (as well as CP2) to the LeuAC urzyme. That consequence is a substantial increase in transition-state stabilization for tRNA^Leu^ aminoacylation by enforcing coupled behavior on the two catalytic signatures, as discussed further in §2.5 and §3.

### 2.5 Nonpolar sidechains V51 and M651 anchor both HVGH and KMSKS signatures securely into the ABD

It is worth noting that although only minimal structural data are available for any aaRS urzymes(35), neither of the hydrophobic residues in the two signature sequences appears from crystal structures of full-length Class I aaRS to be engaged in any significant nonpolar packing interactions in LeuAC itself (Fig. 5C). By contrast, a well-developed packing network of nonpolar amino acids including the two hydrophobic side chains together with residues from the ABD in full-length LeuRS anchors the valine and methionine residues within that domain (Fig. 5). Delaunay tessellation and likelihood scoring(36,37) identified a homologous network in TrpRS (38), and was used here to identify residues detailed in Figure 5D. Similar networks can be identified in all full-length Class I aaRS. They appear both necessary and sufficient to coordinate the behavior of the two catalytic signatures, as shown schematically in Fig. 5E.

**Figure 5.**
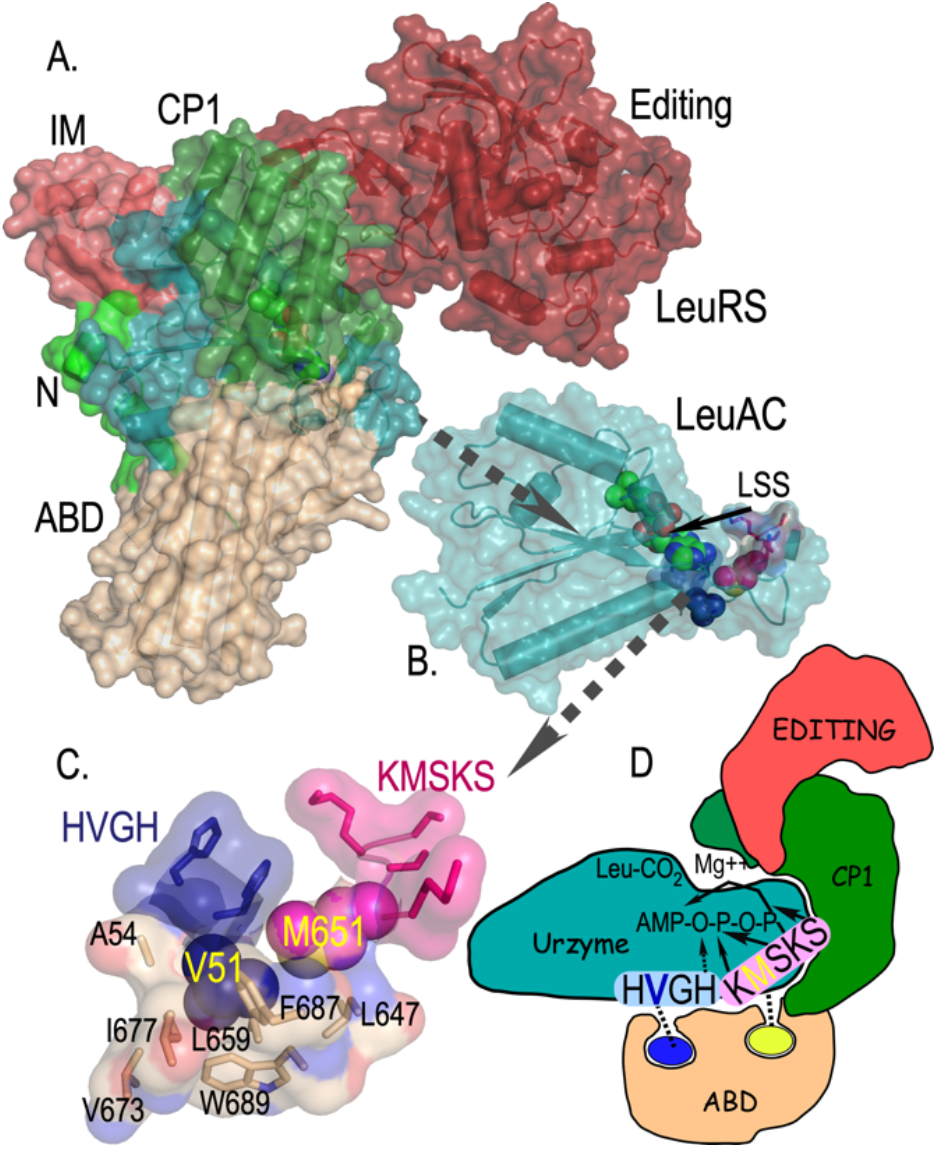
Internal mechanics of the LeuRS (B) and LeuAC (C) active sites. **A**. Space-filling cartoon of intact LeuRS showing the same domains with the same color scheme as in A. **B**. LeuAC urzyme, LSS is a leucyl-5’sufoamyl adenosine analog of the activated amino acyl adenylate. **C**. Cartoon of the nonpolar side chain packing anchoring V51 (blue) from the HVGH (blue) signature and M651 (magenta) from the KMSKS signature firmly within the ABD. Residues that participate in the high statistical potential Delaunay tetrahedra surrounding the nonpolar side chains of the two signature sequences are indicated by sticks and labeled. **D**. Cartoon highlighting the tight mechanical linkage of the two signature sequences to the ABD, and consequently to their cooperative motions relative to the urzyme.

## 3. DISCUSSION

The role of cooperative interactions between active-site residues in enzyme catalysis is a recurring question that is seldom addressed directly by experimentation(39), despite well-established protocols for thermodynamic cycle analysis of combinatorial mutants(34). The thermodynamic cycle is a revealing linear transformation of the experimental transition state stabilization free energies, ΔG^‡^, which as noted elsewhere are additive. The alternative coordinate system highlights the magnitude and sign of individual and energetically coupled contributions to catalysis.

### 3.1 Emergent coupling deepens evidence for the authenticity of urzyme catalysis

We cannot overlook the obvious conclusion that the contrast between the uncoupled behavior of the two catalytic signatures in LeuAC and their strong coupling by the ABD and CP in full length LeuRS reinforces the authenticity of both LeuAC(12) and TrpAC(26) catalysis. The evolution of intramolecular coupling shown here therefore materially strengthens the argument that aaRS urzymes represent valid experimental models for the ancestral assignment catalysts that originally enabled nature to mine the immense functional diversity represented by proteins(7).

### 3.2 The LeuAC studies complement and extend earlier thermodynamic cycle analyses of TrpRS

Class I aminoacyl-tRNA synthetase enzymes (aaRS) afford several examples of how thermodynamic cycle analysis probes structure-function relationships. First and Fersht showed that the effects of mutating residues in the KMSKS loop were localized quite specifically along the reaction profile to destabilize the ground state pre-transition state and stabilize the transition state for tyrosine activation by TyrRS(21). A packing motif(40) called the D1 switch(38) links the two β-strands and α-helix of the first crossover connection to the N-terminal β-strand of the second crossover connection of the Rossmann dinucleotide-binding domain, creating a conformational transition state that mediates shear during relative domain motion of the ABD and CP and imposing discrete pre-transition and products conformations in TrpRS(41-43) and likely in other Class I aaRS.

Combinatorial mutagenesis of four residues from the D1 switch, assayed with both Mg^++^ and Mn^++^ established a five-dimensional thermodynamic cycle. That cycle revealed that the packing motif, which is ∼20 Å from the active-site metal, is nevertheless coupled by –6.2 kcal/mole to the catalytic function of the metal(27). That contribution nearly equals the full catalytic contribution, – 6.4 kcal/mole, of the metal measured by assaying metal-free TrpRS(44). A complementary, modular thermodynamic cycle consisting of TrpRS, its urzyme, TrpAC, and TrpAC plus either the CP insert (i.e. the intact catalytic domain) or the ABD showed that both CP and the ABD alone actually reduced the activity of TrpAC in aminoacylation of tRNA^Trp^, but that both together restored full activity, a contribution of ∼5 kcal/mole—nearly the entire contribution of Mg^++^.(26) Finally, both the combinatorial mutagenesis(25) and modular cycles(26) also implicate the corresponding energetic coupling in amino acid specificity.

### 3.3 Data in Figures 2-4 extend experimental exploration into the time domain(45-52)

The fragmentary vignettes—that the KMSKS signature breaks the synergistic binding of amino acid and ATP in the pre-transition state, the comparable coupling energies between D1 switch residues and Mg^++^ and between the TrpRS ABD and CP domains, and the minimum action path connecting successive structures along the structural reaction profile—consistently suggest that a central function of the ABD and CP domains is to impose coordinated motion on the disparate components of the active site, such that they all assemble simultaneously into a catalytically active, discriminating configuration.

We extend the thermodynamic cycle analysis in this work to the evolutionary time domain to identify quantitatively the explicit gain of function induced in Class I leucyl-tRNA synthetase by the acquisition of the connecting peptide 1 (CP) insertion and anticodon-binding (ABD) domains. The new data reported here reinforce the picture outlined in the previous paragraph. In LeuAC, the absence of the two larger domains leaves the two catalytic signatures uncoordinated, contributing in unexpected ways to transition state stabilization. In full length LeuRS, on the other hand, although neither signature alone can enhance transition-state stabilization, their combination, in the context of the two additional domains, furnishes substantial catalytically productive synergy. That we now have identified much the same phenomena in one of the largest as well as the smallest, Class I aaRS suggests generality in the superfamily. The generality is reinforced by the superfamily-wide conservation of the “enforcer” packing motif illustrated in Fig. 5D, in which high SNAPP potential(36,37) hydrophobic clusters anchor the sole hydrophobic side chains of each catalytic signature, coordinating their motion.

### 3.4 Acquired CP and anticodon-binding domains may have had distinct selective advantages

We have argued that aaRS of both Classes began as 46-residue polypeptides called “protozymes” that accelerated the rate of amino acid activation ∼10^6^-fold(53). That hypothesis gained support from validation by Tamura’s group(54). Thus, it is likely that the HVGH signature has its roots in the protozyme. Class I aaRS urzymes, however, are more than twice as big (∼130 residues), and the KMSKS signature must have roots in that subsequent stage. The selective advantage of the KMSKS signature likely contributed to the acquisition of the second half of urzymes, which accelerate the rate of activation by 10^9^-fold.

As noted previously(26), neither CP nor the ABD enhanced either specificity or catalytic rate enhancement of the TrpRS urzyme. This prompts the question of how either domain might have been selected except in the unlikely case that the urzyme acquired both simultaneously. The AST assays performed to assess the fraction of active catalysts in each sample suggest a possible resolution of that conundrum. The proportion of total ATP usefully converted to AMP production by LeuAC (0.21) is roughly half that (0.4) observed for LeuRS. If the CP domain stabilizes closed, productive forms of the urzyme so that it used ATP more efficiently, that would certainly have given it a selective advantage, even without imposing the coupling between the two signatures. This hypothesis is experimentally testable.

### 3.5 Significant rate acceleration by early polypeptide catalysts did not require sophisticated amino acid side chains

The LeuAC AVGA variant (<ΔG^‡^> = 1.9 kcal/mole) is quantitatively more active than the same variant of the full-length LeuRS (<ΔG^‡^> = 2.1 kcal/mole). That brings us to what is perhaps the most consequential implication from the data in Figures 2-4. The overall rate enhancements observed for amino acid activation by aaRS urzymes excerpted from two Class I (TrpRS(6,7,15), LeuRS(12)) and one Class II aaRS (HisR(10,26)) represent ∼ 60% of the overall transition-state stabilization free energy observed for the corresponding full length aaRS. How much of that must be attributed to the precise orientation of specific amino acid side chains in the active site?

Our data suggest that the surprising answer to that question is: apparently very little, if any of it. Indeed, the superiority of the AVGA LeuAC mutant actually means that the two histidine residues in that highly conserved signature actually reduce the rate of aminoacylation unless they are coupled to the KMSKS sequence by the additional domains. Moreover, the AMSAS mutant is very nearly as active as the WT LeuAC. These observations are consistent with the previous mutational analysis of the TrpAC urzyme, where mutation of an aspartate at the N-terminus of the C-terminal α-helix of the second crossover connection in the Rossmann fold reduces activity by ∼200 fold in TyrRS but increases activity ∼25-fold in the TrpRS urzyme.

We previously noted that secondary structural; i.e., backbone interactions in Class I and II urzymes could account for tRNA(55,56) and amino acid(57) substrate selection(7). Together with our previous evidence that the TrpAC urzyme may be a catalytically active molten globule,(35) these results all substantiate the possibility that substantial catalytic rate enhancement does not require precise orientation of specialized amino acid side chains. It now seems that primordial protein catalysts achieved high rate enhancements and a modicum of substrate specificity largely via physical properties of the active-site pocket that depend largely on backbone interactions that do not require specific amino acid side chains.

That conclusion lends a strong experimental basis for Wong’s coevolution theory for the emergence of the genetic code and it coevolution with metabolic pathways necessary to synthesize amino acids not produced by prebiotic chemistry.(17,18)

## 4. MATERIALS AND METHODS

### 4.1 Expression and purification of LeuRS and LeuAC

The gene for *Pyrococcus horikoshii* (Ph) LeuRS was synthesized by Gene Universal and expressed from pET-11a in BL21-CodonPlus (DE3)-RIPL (Agilent). Cells were grown at 37°C and induced with 300 µM IPTG for 4 hours then harvested and stored overnight at -20°C. The cell pellet was resuspended in 1x Ni-NTA buffer (20 mM Tris, pH 8.0, 300 mM NaCl, 10 mM imidazole, 5 mM β-ME) plus cOmplete protease inhibitor (Roche) and lysed by three 15K psi passes on an Avestin Emulsiflex. Cell debris was pelleted at 4°C 30 minutes 20K rpm. The soluble fraction was heated at 80°C for 30 minutes to denature native *Escherichia coli* proteins. The heated cell extract was then pelleted, and the soluble material was loaded on to an equilibrated Ni-NTA column. The column was washed with three volumes 1x Ni-NTA buffer, then protein was eluted in a stepwise fashion with imidazole concentrations of 40, 80, 100, 200, and 500 mM imidazole. The fractions containing the protein of interest were pooled and dialyzed overnight against 200 mM HEPES, pH 7.4, 450 mM NaCl, 100 mM KCl, 10 mM β-ME. The following day the dialyzed protein was concentrated and mix to 50% glycerol and stored at -20°C.

LeuAC was expressed as an MBP fusion from pMAL-c2x in BL21Star(DE3) (Invitrogen). Cells were grown, induced, harvested, and lysed similarly to Ph LeuRS with the distinct difference of being resuspended in buffer (20 mM Tris, pH 7.4, 1 mM EDTA, 5 mM β-ME, 17.5% Glycerol, 0.1% NP40, 33 mM (NH_4_)_2_SO_4_, 1.25% Glycine, 300 mM Guanidine Hydrochloride) plus cOmplete protease inhibitor (Roche). LeuAC crude extract was then pelleted at 4°C 30 minutes 15K rpm to remove insoluble material. The extract was then diluted 1:4 with Optimal Buffer and loaded onto equilibrated Amylose FF resin (Cytiva). The resin was washed with five column volumes of buffer and the protein was eluted with 10 mM maltose in Optimal Buffer. Fractions containing protein were concentrated and mixed to 50% glycerol and stored at -20°C. All protein concentrations were determined using the Pierce™ Detergent-Compatible Bradford Assay Kit (Thermo Scientific). Experimental assays were performed either with the intact MBP-LeuAC fusion protein or with samples cleaved by tobacco etch virus (TEV) protease, purified as described (58). Purity and cleavage efficiency was determined by running samples on PROTEAN® TGX (Bio-RAD) gels. Some experiments used a second LeuAC variant, from a more recent Rosetta design algorithm in which we attempted to modify surface residues to increase solubility (Matt Cummins, unpublished). Amino acid sequences for both variants are given in the Supplemental data and compared to the native Ph LeuRS sequence in Supplemental Figure S1. We could not detect any significant differences between results from the two different LeuAC fusion constructs.

### 4.2 Single turnover active-site titration assays

Active-site titration assays were performed as described (29,59). 3 µM of protein was added to 1x reaction mix (50 mM HEPES, pH 7.5, 10 mM MgCl_2_, 5 µM ATP, 50 mM amino acid, 1 mM DTT, inorganic pyrophosphatase, and α-labeled [^32^P] ATP) to start the reaction. The α labeling position allowed us to follow time courses ADP and AMP production, as well as for ATP consumption. Timepoints were quenched in 0.4 M sodium acetate 0.1% SDS and kept on ice until all points had been collected. Quenched samples were spotted on TLC plates, developed in 850 mM Tris, pH 8.0, dried and then exposed for varying amounts of time to a phosphor image screen and visualized with a Typhoon Scanner (Cytiva). The ImageJ measure function was used to quantitate intensities of each nucleotide. The time-dependent of loss (ATP) or de novo appearance (ADP, AMP) of the three adenine nucleotide phosphates were fitted using the nonlinear regression module of JMP16PRO™ to equation (1):

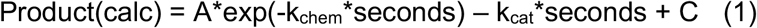

where k_chem_ is the first-order rate constant, k_cat_ is the rate of turnover, A is the amplitude of the first-order process, and C is an offset. Eqn. (1) was also used for convenience to fit the appearance of nucleotide products, AMP, and ADP. In that case the roles of A and C are inverted, C being the burst size and A the offset and the signs of both rate constants are given the opposite sign.

#### Burst size estimation from fitted parameters in Eqn. (1)

For [ATP] decay curves, the fitted A value estimates n directly as n = A*[ATP]/[Enzyme]. C gives n =(1–C)**[ATP]/[Enzyme. For exponentially increasing concentrations of product, the situation reverses, and n = (1–A) *[ATP]/[Enzyme and n = C*[ATP]/[Enzyme. These approximations are justified by the very small variance of multiple estimates, and by the internal self-consistency discussed in §

### 4.2 tRNA^Leu^ aminoacylation assays

A plasmid encoding the *P. horikoshii* tRNA^Leu^ (TAG codon) was synthesized by Integrated DNA Technologies and used as template for PCR amplification of the tRNA and upstream T7 promoter and downstream Hepatitis Delta Virus (HDV) ribozyme. The PCR product was used directly as template for T7 transcription. Following a 4-hour transcription at 37°C the RNA was cycled five times (90°C for 1 min, 60°C for 2 min, 25°C for 2 min) to increase the cleavage by HDV. The tRNA was purified by urea PAGE and crush and soak extraction. The tRNA 2’-3’ cyclic phosphate was removed by treatment with T4 PNK (New England Biolabs) following the manufacturer’s protocol. The tRNA was then phenol chloroform isoamyl alcohol extracted, filter concentrated, aliquoted, and stored at -20°C.

We determined the active fraction of tRNA^Leu^ by following extended acylation assays using full-length *P. horikoshii* LeuRS until they reached a plateau. That plateau value was 0.38, which we used to compute tRNA^Leu^ concentrations in assays with both LeuRS and LeuAC.

Aminoacylations were performed in 50 mM HEPES, pH 7.5, 10 mM MgCl_2_, 20 mM KCl, 5 mM DTT with indicated amounts of ATP and amino acids. Desired amounts of unlabeled tRNA— mixed with [α^32^P] A76-labeled tRNA for assays by LeuAC— were heated in 30 mM HEPES, pH 7.5, 30 mM KCl to 90°C for 2 minutes. The tRNA was then cooled linearly (drop 1°C/30 seconds) until it reached 80°C when MgCl_2_ was added to a final concentration of 10 mM. The tRNA continued to cool linearly until it reached 20°C.

### 4.4 Data processing and Statistical analysis

Phosphorimaging screens of TLC plates were densitometered using ImageJ. Data were transferred to JMP16PRO™ Pro 16 via Microsoft Excel (version 16.49), after intermediate calculations. We fitted active-site titration curves to Eqn. (1) using the JMP16PRO™ nonlinear fitting module. R2 values were all > 0.97 and most were > 0.99.

Factorial design matrices in Tables 1,2 were processed using the Fit model multiple regression analysis module of JMP16PRO™ Pro, using an appropriate form of equation (2)

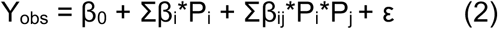

where Y_obs_ is a dependent variable, usually an experimental observation, β_0_ is a constant derived from the average value of Y_obs_, β_i_ and β_ij_ are coefficients to be fitted, P_i,j_ are independent predictor variables from the design matrix, and ε is a residual to be minimized. All rates were converted to free energies of activation, ΔG^‡^ = -– RTln(k), before regression analysis because free energies are additive, whereas rates are multiplicative. For example, the activation free energy for the first-order decay rate in single-turnover experiments is ΔG^‡^k_chem_.

Multiple regression analyses of factorial designs exploit the replication inherent in the full collection of experiments to estimate experimental variances on the basis of t-test P-values, in contrast to the presenting error bars showing the variance of individual datapoints. Multiple regression analyses reported here also entail triplet experimental replicates, which enhance the associated analysis of variance.

## Data availability

All data (The Protein Databank) or upon request from the author.

## Author contributions

J.H. and Q.C.T. designed the combinatorial LeuAC mutants and performed all assays. Q.C.T. constructed and assayed the combinatorial LeuRS mutants. C.W.C., J.H., and G.C.T. all performed data analysis. J.D. provided phylogenetic insight for Figure 1s as well as the evolutionary history of the signature sequences. C.W.C. wrote the manuscript, and all authors contributed to and approved the final figures and text.

## Acknowledgments

P. Wills provided continual constructive discussions. E. First and L. Betts made helpful comments on the manuscript.

